# Endoplasmic reticulum stress activates human IRE1α through reversible assembly of inactive dimers into small oligomers

**DOI:** 10.1101/2021.09.29.462487

**Authors:** Vladislav Belyy, Iratxe Zuazo-Gaztelu, Andrew Alamban, Avi Ashkenazi, Peter Walter

## Abstract

Protein folding homeostasis in the endoplasmic reticulum (ER) is regulated by a signaling network, termed the unfolded protein response (UPR). Inositol-requiring enzyme 1 (IRE1) is an ER membrane-resident kinase/RNase that mediates signal transmission in the most evolutionarily conserved branch of the UPR. Dimerization and/or higher-order oligomerization of IRE1 are thought to be important for its activation mechanism, yet the actual oligomeric states of inactive, active, and attenuated mammalian IRE1 complexes remained unknown. We developed an automated two-color single-molecule tracking approach to dissect the oligomerization of tagged endogenous human IRE1 in live cells. In contrast to previous models, our data indicate that IRE1 exists as a constitutive homodimer at baseline and assembles into small oligomers upon ER stress. We demonstrate that the formation of inactive dimers and stress-dependent oligomers is fully governed by IRE1’s lumenal domain. Phosphorylation of IRE1’s kinase domain occurs more slowly than oligomerization and is retained after oligomers disassemble back into dimers. Our findings suggest that assembly of IRE1 dimers into larger oligomers specifically enables *trans-* autophosphorylation, which in turn drives IRE1’s RNase activity.

## Introduction

Protein oligomerization is central to cell biology. The regulated assembly of membrane proteins into dimers or larger oligomers constitutes a fundamental cellular mechanism for relaying information across membranes. The majority of receptor superfamilies rely on some form of oligomerization, including G-protein coupled receptors (*1*), integrins (*2*), receptor tyrosine kinases (*3*), T-cell receptors (*4*) and death receptors (*5*). While significant progress has been made in understanding oligomeric assembly of cell-surface receptors, much less is known about oligomerization of intracellular membrane proteins. This is in part because intracellular oligomers are often too small, dynamic, or weakly associated to be resolved by conventional approaches.

One such oligomer-forming protein is the ER membrane-resident stress sensor IRE1. It is a dual-function kinase/ribonuclease (RNase) responsible for initiating the most evolutionarily conserved branch of the unfolded protein response (UPR) (*6*–*8*). The UPR is a major signaling network that lies at the core of cellular homeostasis and is responsible for making cellular life-or-death decisions when faced with an imbalance between load on and capacity of the ER’s protein folding machinery (*9*, *10*). IRE1’s role as a master regulator of the UPR has made it an important subject of both basic and translational investigation. Upon its activation by the buildup of unfolded proteins in the ER lumen, IRE1 undergoes kinase-mediated *trans*-autophosphorylation and catalyzes RNase-mediated non-conventional splicing of the *XBP1* mRNA (*11*, *12*) (in mammals; *HAC1* mRNA in yeast) (*13*) as well as the decay of multiple mRNA targets (*14*–*16*). While IRE1’s activation is generally thought to involve the formation of dimers and/or larger oligomers, the extent and functional importance of this oligomerization phenomenon, along with the precise oligomeric state of IRE1 complexes, remain hotly debated.

Early work on yeast IRE1 revealed that both the lumenal (*17*) and cytosolic (*18*) domains individually crystallize as helical filaments, and that IRE1 molecules assemble into puncta in the ER membrane upon induction of ER stress (*19*). Similarly, fluorescently tagged human IRE1α (“IRE1” hereafter) was observed to reversibly assemble into large, topologically complex puncta in a stress-dependent fashion (*20*–*22*). The Hill coefficients for purified yeast (*18*) and human (*20*) IRE1 kinase/RNase domains were measured to be ~8 and ~3.4, respectively, indicating that the cooperative formation of oligomers larger than dimers plays an important role in IRE1’s enzymatic cycle. The lumenal domain, while itself lacking catalytic activity, was also observed to assemble into dimers and larger oligomers *in vitro* (*23*, *24*). This assembly occurs across two predicted interfaces: IF1^L^ (L for lumenal), generally accepted to be the primary dimerization interface, and IF2^L^ (**Fig. 1A**), which mediates higher-order oligomerization.

**Figure 1:**
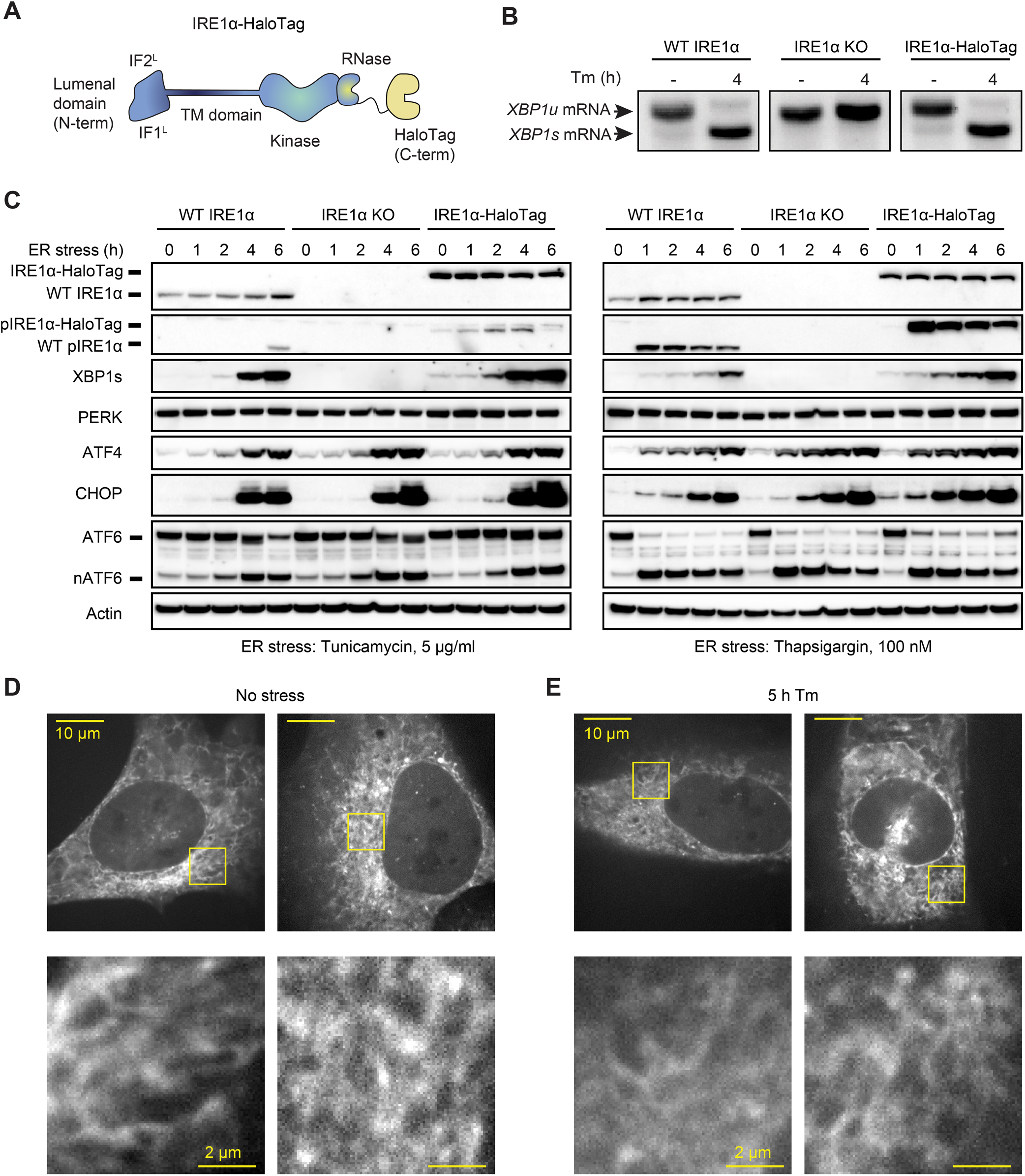
Endogenously tagged IRE1α is fully active despite not forming large clusters. (A) Schematic representation of IRE1 with a C-terminal HaloTag, the construct used for tagging IRE1 at the endogenous locus. IF1L and IF2L refer to the primary dimerization and oligomerization interfaces of the lumenal domain, respectively. (B) RT-PCR analysis of stress-dependent XBP1 mRNA splicing in WT U-2 OS cells, IRE1 knock-out (KO) U-2 OS cells, and U-2 OS cells in which IRE1 has been fully edited with a C-terminal HaloTag. Tm indicates treatment with 5 μg/ml tunicamycin. (C) Immunoblot of UPR activation in response to 5 μg /ml tunicamycin (left) and 100 nM thapsigargin (right) treatments in the three cell lines shown in panel B. (D) Maximum intensity projections of representative spinning-disk confocal images of live cells expressing endogenously tagged IRE1-HaloTag, labeled with the JF549 dye. Regions shown with yellow boxes are enlarged below. (E) Same as D, except the cells have been treated with 5 μg/ml tunicamycin for 5 hours.

Despite this wealth of information, the oligomeric state of both active and inactive IRE1 complexes in mammalian cells remains unclear. It has been alternatively proposed that the monomer-to-dimer transition serves as the main activation signal and that the formation of high-order oligomers is instead the primary regulatory step. The former is supported by the observation of stress-induced increase in crosslinking of a Q105C mutant engineered into the IF1^L^ interface (*25*), while the latter rests on the observation of large clusters of fluorescently tagged IRE1 in stressed cells and on the finding that genetic disruption of the IF2^L^ interface abrogates IRE1 activity (*24*). However, crosslinking of a single residue is not necessarily proportional to the degree of dimerization. Indeed, the dimer of IRE1’s lumenal domains has been predicted to undergo substantial conformation changes upon peptide binding (*24*), which alongside the biochemical changes in the lumen of an acutely stressed ER may alter crosslinking efficiency. Most other studies relied on exogenous overexpression of tagged IRE1, which may in turn bias the equilibrium of an oligomerization-prone protein away from physiologically relevant levels. To pursue an orthogonal strategy, we set out to directly measure the oligomerization of endogenously labeled IRE1 in live human cells. To this end, we developed a single-molecule microscopy approach that proved useful to reveal the precise oligomeric changes that underpin IRE1 activation. More broadly, this approach promises to provide a powerful tool to study the oligomerization of other proteins residing on internal membranes in eukaryotic cells.

## Results

### Endogenously tagged IRE1 is fully active despite not forming large clusters

To study the oligomerization of endogenous IRE1, we inserted a C-terminal HaloTag (*26*) into IRE1’s genomic locus in U-2 OS cells using CRISPR/Cas9-based gene editing (**Fig. 1A**). Following clonal selection, we chose a clone that satisfied the following criteria: 1) comparable IRE1 expression levels to unedited U-2 OS cells, 2) absence of wild-type (WT) IRE1 protein lacking the HaloTag, and 3) intact UPR activation in response to ER stress. The latter was ascertained by the detection of ER stress-dependent *XBP1* mRNA splicing (**Fig. 1B**), IRE1 phosphorylation, production of XBP1s protein, upregulation of the CHOP and ATF4 transcription factors, and cleavage of ATF6 (**Fig. 1C**). We concluded that the endogenous C-terminal HaloTag does not substantially interfere with IRE1’s kinase or RNase activity and could provide an excellent way to image IRE1 dynamics in live cells.

A key advantage of HaloTag fusion proteins stems from the fact that they can be labeled with bright and photostable cell-permeable dyes. Thus, despite IRE1 being a comparatively low-abundance protein, we could readily image it by spinning-disk confocal microscopy after labeling it with the JF549 dye conjugated with the HaloTag ligand (*27*). As expected, IRE1-HaloTag exhibited a reticulated distribution characteristic of ER-localized proteins (**Fig. 1D**). We were surprised to observe a total lack of detectable cluster formation upon induction of ER stress (**Fig. 1E**), in direct contrast to previous work by us and others that relied on ectopic expression of GFP-tagged IRE1 protein (*20*, *21*, *28*–*32*). While unexpected, this observation does not rule out lower-order IRE1 oligomerization at endogenous expression levels, since the limited sensitivity of confocal microscopy would preclude the detection of small oligomers such as dimers or tetramers as distinct morphological features. We therefore sought to devise a more sensitive approach for detecting small oligomers in the ER membrane.

### Development of a two-color tracking algorithm for the detection of small oligomers

Detection of small protein oligomers inside intact cells is a notoriously challenging task. A range of approaches, each carrying a unique set of strengths and limitations, had been employed in the past (*33*–*36*). We leveraged the fact that the HaloTag protein can be labeled with fluorophores of different colors.

In principle, if a protein is stochastically labeled by fluorophores with distinct spectra and subsequently imaged with single-molecule resolution, its average oligomeric state can be determined by quantifying the fraction of particles that fluoresce in more than one color (**Fig. 2A, B**). However, to date this approach has been limited to reconstituted *in vitro* systems and plasma membrane-bound proteins (*37*–*44*). Furthermore, previous implementations lacked experimental controls of defined stoichiometry, relying on a number of physical assumptions to estimate the degree of oligomerization. To overcome these challenges, we developed a fully automated image analysis pipeline for identifying co-localizing two-color trajectories of ER membrane-resident proteins.

**Figure 2:**
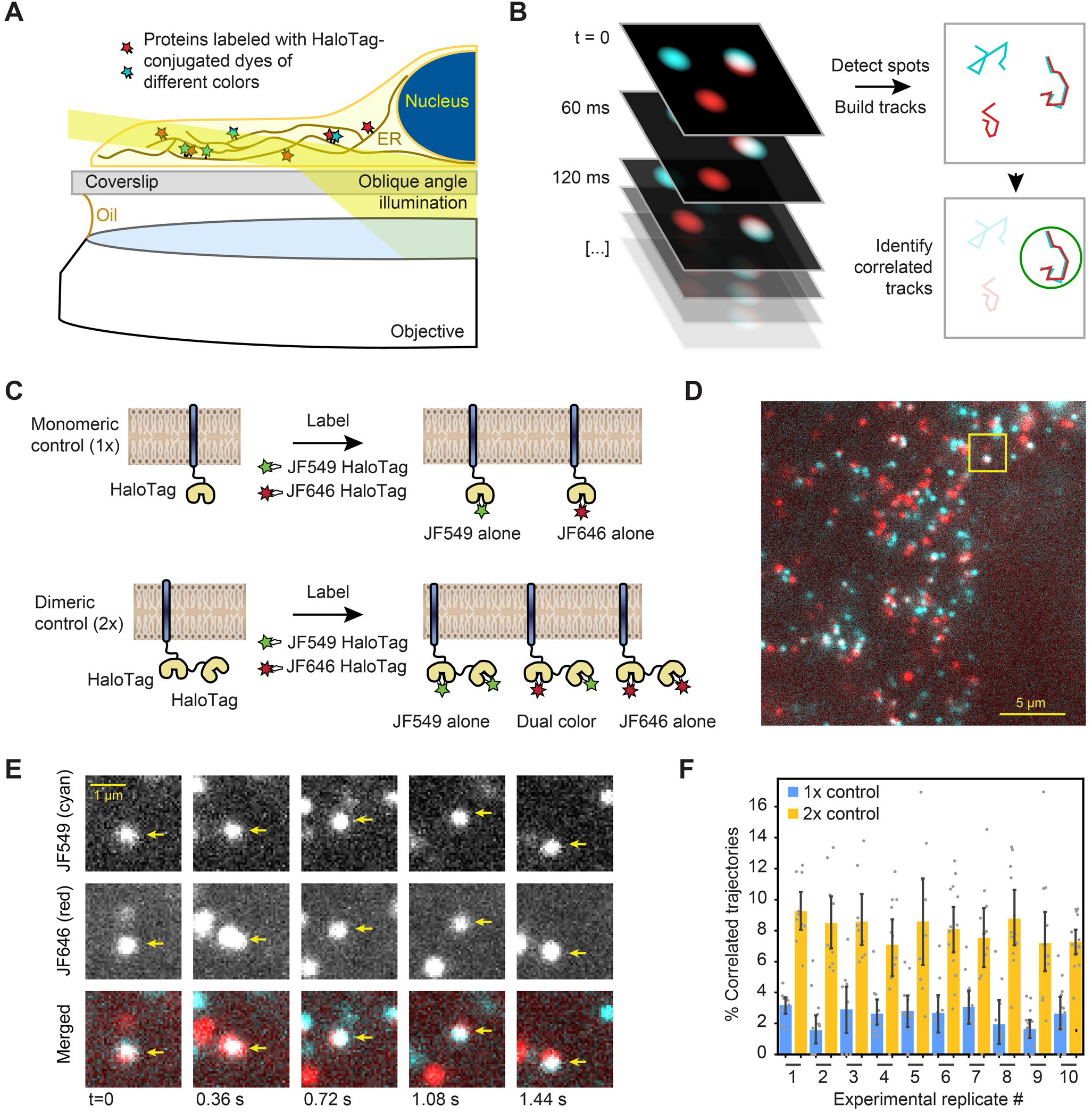
Single-particle tracking approach for detection of small oligomers. (A) Schematic depiction of the assay. Cells expressing low levels of HaloTag-conjugated proteins are labeled with a mixture of HaloTag-conjugated dyes and imaged by oblique angle illumination. (B) Principle behind the analysis of single-particle data. Fluorescent spots are independently tracked in two channels, and correlated trajectories are identified computationally. (C) Design of the 1x and 2x HaloTag controls. (D) Representative frame from a movie of a cell expressing an ER-tethered 2x tandem HaloTag and labeled with a mixture of JF549 (cyan) and JF646 (red) dyes. (E) Several frames of the boxed region in panel D, with co-localizing spots identified with arrows. (F) Percentage of correlated trajectories from cells expressing the 1x and 2x HaloTag controls, comparing data collected in ten independent experimental replicates. Each data point represents a single cell, typically comprising several hundred trajectories. Error bars represent 95% confidence intervals.

First, we calibrated our tracking-based approach using ER membrane-tethered proteins with well-defined oligomeric states. We expressed in U-2 OS cells synthetic constructs containing either a single ER-targeted HaloTag or two HaloTags in tandem (**Fig. 2C**), under control of a weakened CMVd3 promoter (*45*). After labeling to saturation with a mixture of JF-549 HaloTag and JF-646 HaloTag dyes, we imaged the cells by oblique angle illumination microscopy. In longer-exposure movies, it was apparent that both single and double HaloTag constructs exhibited a reticulated distribution characteristic of the ER. The thin, spread-out morphology of U-2 OS cells, together with the exceptional photophysical properties of the JF dyes, allowed us to readily distinguish single diffusing molecules in both channels and track them over multiple frames (**Fig. 2D, E**). As expected, a large number of seconds-long correlated two-color trajectories were observed in cells expressing the tandem 2x HaloTag construct (**Fig. 2D**, **Supp. Movie 1**), but not in cells expressing the single HaloTag construct.

To quantify the fraction of co-localizing spots, we employed the following algorithm. First, spots were automatically detected and tracked in both channels. Then, the tracks were binned into short trajectories using a sliding window of either 12 or 14 frames (0.72 or 0.84 s) to minimize the ambiguity in assignment of crossing tracks. Pearson’s correlation coefficients were then calculated between both the x- and y-coordinates of adjacent tracks within each sliding window. Every track in the JF-549 channel that contained at least one window with a correlation coefficient above a predetermined threshold was classified as a co-localizer (see **Materials and Methods** for details). By repeating this analysis on data collected from cells expressing the 1x and 2x HaloTag controls in ten independent replicates (>10 cells and >1500 trajectories per condition in each replicate), we verified that the algorithm robustly and reproducibly distinguishes between monomeric and dimeric molecules in the ER membrane (**Fig. 2F**). To rule out the remote possibility that the 2x tandem HaloTag protein may be an imperfect control due to differential ligand accessibility of the internal and C-terminal HaloTag proteins, we repeated the analysis with a construct that instead relies on GST dimerization to bring two ER-bound C-terminal HaloTag proteins together. The measured percentage of co-localized trajectories for this construct was statistically indistinguishable (p = 0.50, two-tailed T-test) from that of the tandem 2x HaloTag protein (**Supp. Fig. 1**), thus confirming our ability to unambiguously discern monomers from dimers in live cells.

### IRE1 transitions from dimers to small oligomers upon ER stress

Having validated the ability to detect even small changes in oligomeric state, we applied our analysis to cells expressing endogenously tagged IRE1 (**Fig. 3A**). We could clearly observe individual fluorescent spots corresponding to single IRE1 molecules moving along ER tubules (**Fig. 3B, C**). A fraction of diffusing spots co-localized between the two channels, indicating a significant degree of IRE1 oligomerization even in the absence of stress induction (**Fig. 3D, E**). In fact, upon quantification, the fraction of co-localized IRE1 trajectories in non-stressed cells appeared nearly identical to that of the double-HaloTag control, strongly suggesting that nearly all IRE1 proteins are pre-assembled into dimers at baseline (**Fig. 3F**). Treatment with the glycosylation inhibitor and potent UPR activator tunicamycin (Tm) resulted in a pronounced increase in the fraction of correlated trajectories after 4 hours, indicating that a significant fraction of IRE1 dimers assembled into higher-order oligomers. A simple combinatoric model estimates that the mean number of molecules per cluster increases from ~1.9 to ~2.7 upon Tm stress (see **Materials and Methods** for details). Since our approach does not reveal the individual oligomeric state of any given tracked protein, this observed change is most readily explained by a Tm-dependent shift in equilibrium towards a mixture of dimers and tetramers. Extending the treatment to 24 hours reversed the shift in correlated trajectories, suggesting that IRE1 oligomers dissociate back into dimers under prolonged stress. This finding parallels the previously observed attenuation of IRE1 activity upon prolonged, unmitigated ER stress (*20*, *21*, *46*, *47*) (**Fig. 3F**). Addition of Tm did not induce an increase in the fraction of correlated trajectories of the 1x and 2x HaloTag controls (**Supp. Fig. 2**), demonstrating that the effect is a *bona fide* feature of IRE1 signaling rather than a consequence of stress-dependent remodeling of the ER membrane.

**Figure 3:**
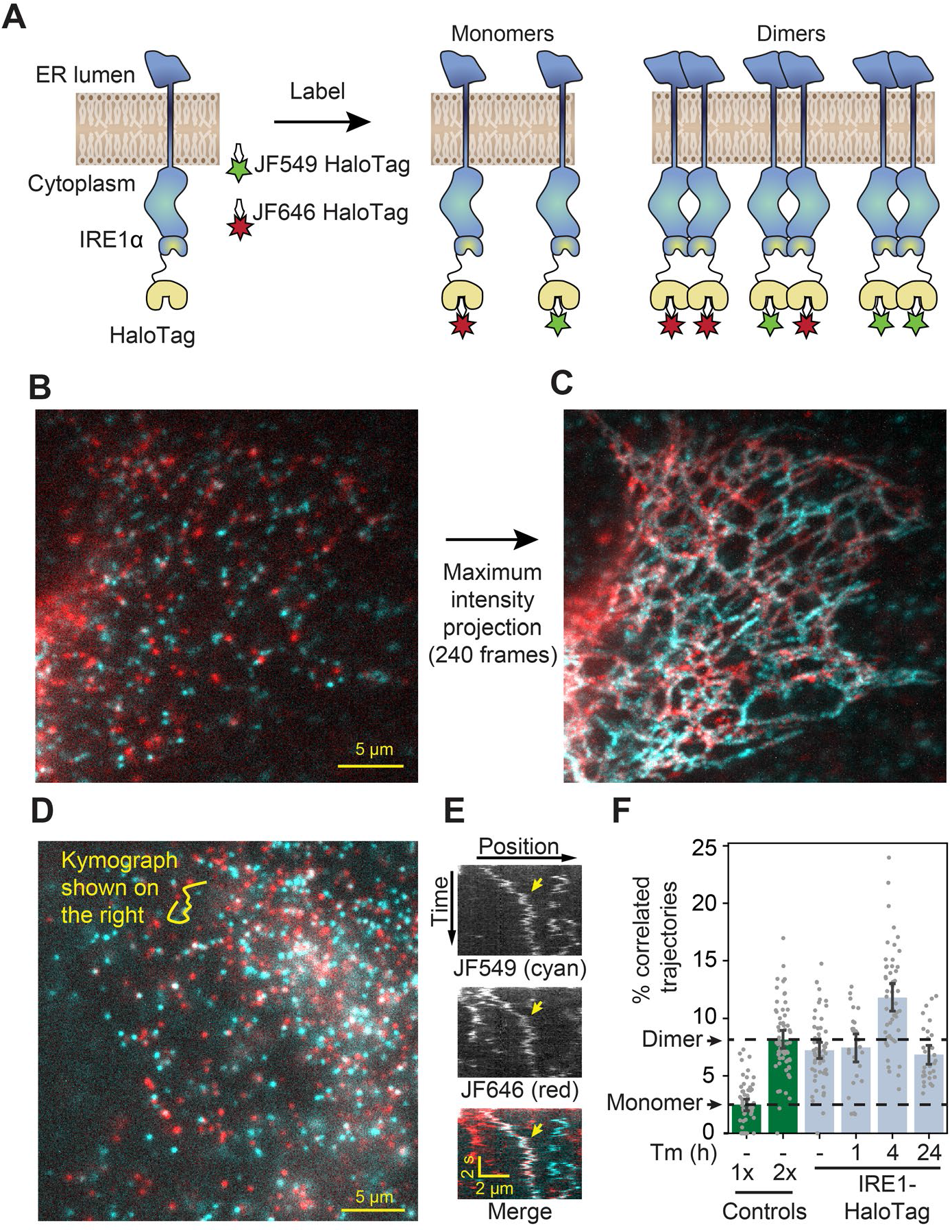
Detection of IRE1 dimers and oligomers in live cells. (A) Schematic depiction of the assay. IRE1-HaloTag is simultaneously labeled with HaloTag dyes of two different colors, JF549 and JF646. If the protein is purely monomeric, all single-molecule tracks are expected to be either one color or the other. If it is purely dimeric, a fraction of tracks will contain both colors. (B) Single frame from a long-exposure movie (100 ms per frame) of a cell in which IRE1-HaloTag is labeled with a mixture of JF549 (cyan) and JF646 (red) dyes. (C) Maximum intensity projection of the entire movie from panel B showing that single IRE1 molecules diffuse along ER tubules. (D) Single frame from a short-exposure movie (50 ms per frame) of a cell in which IRE1-HaloTag is labeled with a mixture of JF549 (cyan) and JF646 (red) dyes. (E) Kymograph (time vs. position plot) along the line shown in panel D. Co-localizing diffusional IRE1 trajectory is shown with a yellow arrow. (F) Stress-induced changes in IRE1 oligomerization in response to treatment with 5 μg/ml tunicamycin (Tm), as quantified by the fraction of correlated trajectories. Green bars on the left correspond to the 1x and 2x HaloTag controls, respectively. Error bars represent 95% confidence intervals.

A key aspect of IRE1 activation is its *trans-*autophosphorylation.

Intriguingly, thapsigargin (Tg), which disrupts ER calcium homeostasis by blocking sarco/endoplasmic reticulum Ca^2+^ pumps (*48*), induced IRE1 phosphorylation much more rapidly and strongly than Tm, despite leading to similar overall levels of XBP1s production (**Fig. 1C**). This observation prompted us to test whether oligomerization is directly proportional to IRE1 phosphorylation by comparing the effects of different ER stressors. Treatment with dithiothreitol (DTT), which causes protein misfolding by reducing disulfide bonds, induced IRE1 oligomerization to the same extent as Tm (**Fig. 4A**). However, to our surprise, treatment with a high concentration (100 nM) of Tg did not induce a detectable change in oligomeric state either 2 or 4 hours after treatment. Since Tg is a fast-acting stressor compared to Tm, we reasoned that the apparent lack of oligomerization in response to Tg might be explained by a rapid formation and dissolution of IRE1 oligomers, which could be effectively complete by the 2-hour time-point. Indeed, imaging cells only 10 minutes after the addition of 100 nM Tg revealed a robust increase in IRE1 oligomerization, as indicated by an increase in the fraction of correlated trajectories (**Fig. 4A**). Furthermore, a lower concentration of 1 nM Tg led to IRE1 oligomerization at the longer 2- and 4-hour time-points. Meanwhile, when 100 nM Tg was combined with saturating Tm, there was no detectable IRE1 oligomerization 4 hours after treatment, demonstrating that the repressive effect of extended Tg treatment overrides the pro-oligomerization effect of Tm. Taken together, our results show that IRE1 phosphorylation lags behind oligomerization and that all commonly used ER stressors induce IRE1 oligomerization, albeit on different temporal scales.

**Figure 4:**
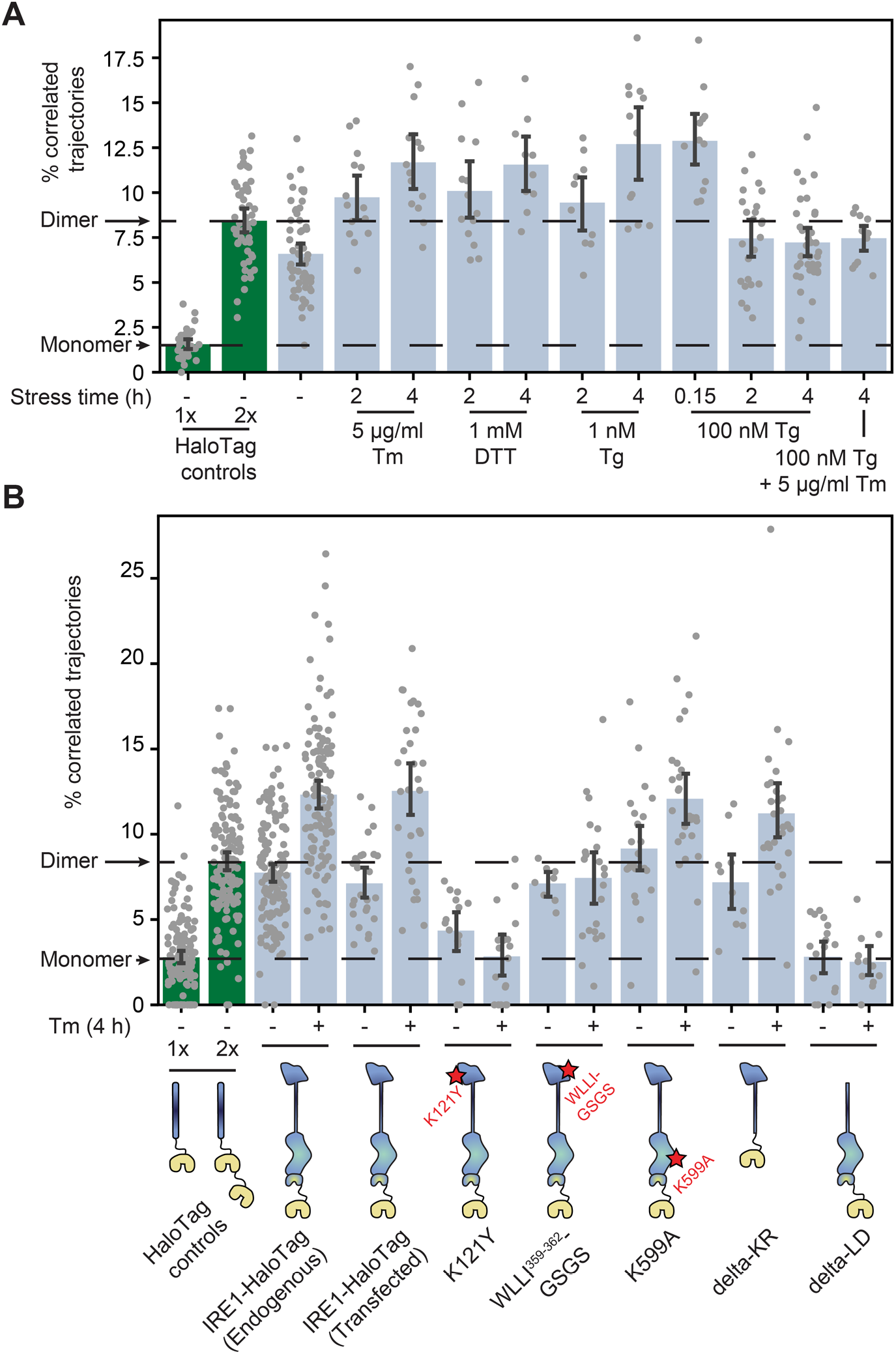
Effects of stressors and mutations on IRE1 oligomerization. (A) Oligomerization of endogenously tagged IRE1-HaloTag in U-2 OS cells treated with the indicated ER stressors for the indicated amounts of time. (B) Oligomerization of the indicated IRE1 mutants transiently transfected into IRE1 KO U-2 OS cells and expressed under the control of the weak CMVd3 promoter. “IRE1-HaloTag (endogenous)” refers to the endogenously tagged IRE1 cells that are shown in panel A. Error bars represent 95% confidence intervals.

Next, we sought to exclude the formal possibility that the observed dimer-to-oligomer transition was either a clonal artifact or a consequence of differences in expression levels of IRE1 and the control constructs. To this end, we used a second clone of IRE1-HaloTag cells that expressed severalfold less IRE1-HaloTag than the amount of WT IRE1 present in parental cells (**Supp. Fig. 3A, B**). This clone exhibited reduced *XBP1* mRNA splicing activity (**Supp. Fig. 3C**), yet our single-particle analysis revealed that IRE1 remained dimeric in unstressed cells even at this decreased expression level (**Supp. Fig. 3D**). Meanwhile, the extent of stress-induced oligomerization was markedly reduced, mirroring the reduction in RNase activity. To specifically address any potential discrepancies between expression levels of IRE1 and the HaloTag controls, we plotted the measured fraction of correlated trajectories against the total number of trajectories in a given movie. The total number of trajectories served as a proxy for the density of fluorescent spots and, by extension, for the relative abundance of the protein in a cell. We found that protein abundance was comparable for IRE1-HaloTag and the HaloTag controls, and that the differences in the fraction of correlated trajectories remained robustly detectable across a wide range of spot densities (**Supp. Fig. 4**). We concluded that both the formation of stable IRE1 dimers in non-stressed cells and the assembly of IRE1 into larger oligomers upon stress induction were neither peculiarities of a single clone nor expression level artifacts, but rather genuine features of IRE1 biology.

### The lumenal domain governs the formation of both dimers and oligomers

To determine which regions of IRE1 are responsible for assembling the protein into dimers and oligomers, we applied our trajectory analysis to a number of key IRE1 mutants (**Fig. 4B**). Since constructing each mutant by CRISPR technology would have been impractical, we first checked whether the dimer-to-oligomer transition could be measured with transiently transfected IRE1-HaloTag. Indeed, when we expressed IRE1-HaloTag under the control of the truncated CMVd3 promoter in IRE1 KO U-2 OS cells, our method unambiguously detected both the presence of IRE1 dimers in the non-stressed cells and a shift towards larger oligomers upon treatment with Tm. This result paved the way for testing IRE1 mutants in a similar fashion. The first revealing pair of functional mutants comprised K121Y, which disrupts the IF1^L^ interface of the IRE1 lumenal domain (*20*), and WLLI^359-362^-GSGS, which disrupts the lumenal domain’s oligomerization interface IF2^L^ (*24*). These two mutations yielded two starkly different outcomes. The oligomeric state of K121Y remained similar to that of the single-HaloTag control, both with and without induction of ER stress. In contrast, WLLI^359-362^-GSGS retained the same oligomeric state as unstressed WT IRE1 both with (p = 0.71) and without (p = 0.18) ER stress. In other words, both lumenal domain mutants lose the ability to change their oligomeric states in the response to ER stress, with K121Y remaining mostly monomeric and WLLI^359-362^-GSGS remaining mostly dimeric.

Next, we probed the potential contribution of the kinase/RNase domain. K599A, a mutation that abrogates IRE1’s kinase activity (*11*), closely mirrored the phenotype of WT IRE1, with only a slightly reduced difference in oligomerization between the non-stress and stress conditions (**Fig. 4B**). We then tested a pair of more radical mutations: delta-KR, a complete deletion of the kinase/RNase domain, and delta-LD, a complete deletion of the lumenal domain. Remarkably, the delta-LD construct remained purely monomeric regardless of ER stress (p = 0.95 and p = 0.62 for unstressed and stressed cells, respectively, when compared to the 1x HaloTag control), while the delta-KR construct recapitulated the stress-dependent transition from dimers to higher-order oligomers (**Fig. 4B**). Particle density analysis confirmed that results from all IRE1 mutants examined are not correlated with the expression levels of the different constructs and represent the mutants’ intrinsic propensities for oligomerization (**Supplementary Fig. 5**). Taken together, the mutant data confirm the previously proposed role of the lumenal domain as the central governor of IRE1 oligomerization, consistent with its role as the ER stress sensor domain. Both constitutive dimerization and the dimer-to-oligomer transition appear to be entirely controlled by the lumenal domain of IRE1, with the kinase and RNase domains acting downstream.

**Figure 5:**
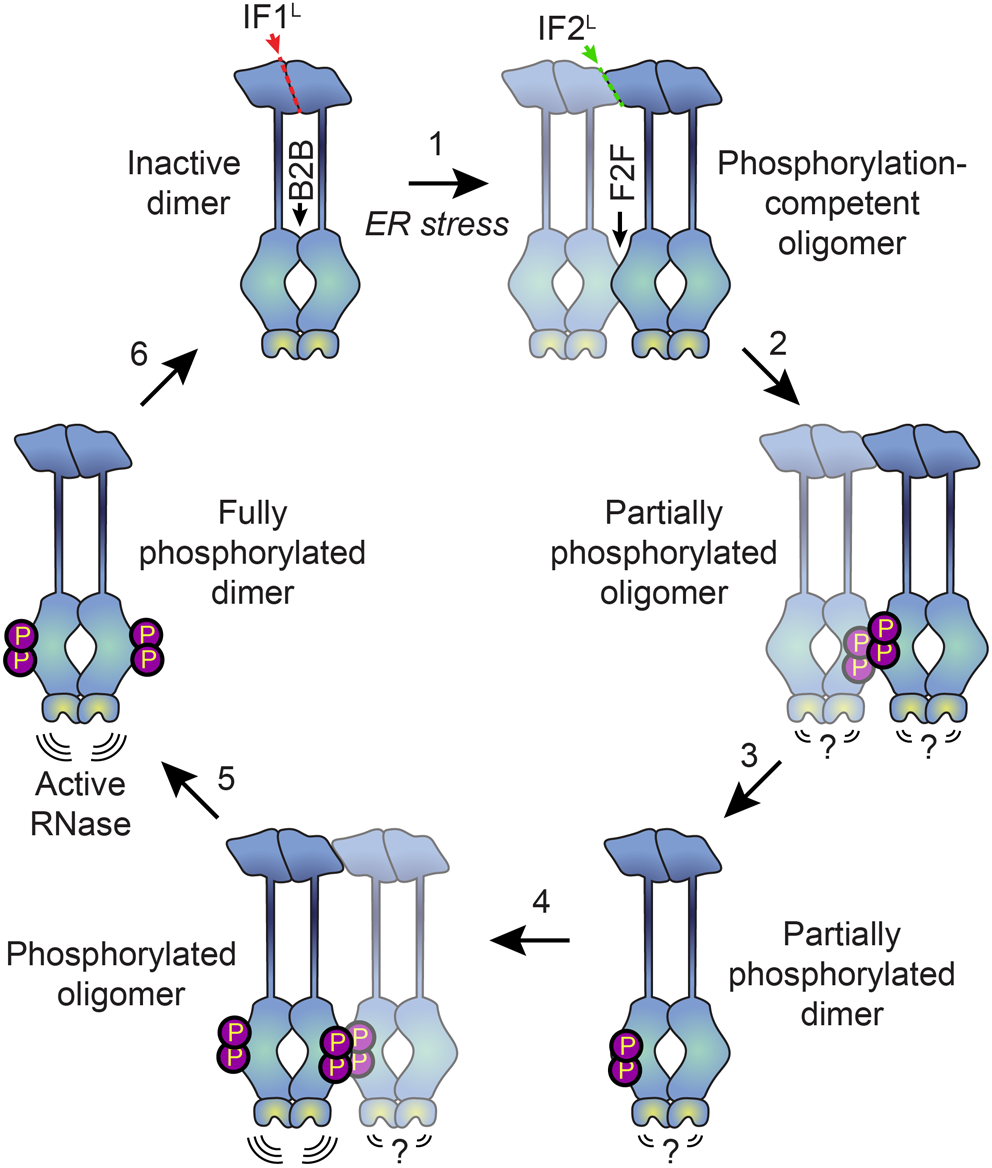
Proposed model for human IRE1 activation. In the absence of external stress, IRE1 is pre-assembled into inactive unphosphorylated dimers via the IF1L interface of the lumenal domain. Kinase domains within the dimer are positioned in a back-to-back (B2B) orientation, which does not allow for phosphorylation. (1) ER stress forces dimers to oligomerize via the IF2L interface, placing the kinase active sites of adjacent dimers in a face-to-face (F2F) orientation that favors *trans-* autophosphorylation. Here and throughout the figure, the original dimer is shown in solid blue tones while the newly associated dimer is semi-transparent. (2) Phosphorylation at the F2F interface results in a partially phosphorylated oligomer, wherein one protomer of each dimer is phosphorylated and one is not. The relative activity of the RNase domains in this state is unknown. (3) At this point, the oligomer may dissociate into partially phosphorylated dimers. (4) Another dimer associates with the partially phosphorylated dimer *via* the second IF2L interface, catalyzing phosphorylation of the second protomer of the original dimer. Note that this may either occur sequentially, as shown here, or simultaneously with step 2, if multiple dimers assemble into a hexamer or larger oligomer. (5) Phosphorylated IRE1 now has an active RNase domain and dissociates into fully active dimers. (6) Eventually, dimers are dephosphorylated by phosphatases and return back into the inactive state.

## Discussion

We measured the stress-dependent oligomeric changes of fully active human IRE1 in live cells at physiological expression levels. Despite IRE1 not forming the massive clusters that were previously observed by us and others in the context of overexpression, we show that the formation of high-order oligomers remains a conserved feature of IRE1’s activation. Surprisingly, IRE1 forms constitutive inactive dimers in the absence of externally induced ER stress in U-2 OS cells, thus challenging the widely held notion (*25*, *49*, *50*) that the monomer-to-dimer equilibrium constitutes the primary regulatory step in IRE1 activation. We demonstrate that the lumenal domain serves as the primary governor of dimerization in the absence of induced stress (via the IF1^L^ interface) as well as oligomer formation in response to ER stress (via the IF2^L^ interface). Indeed, the lumenal domain alone is sufficient for the formation of both resting-state dimers and stress-induced oligomers when tethered to the ER membrane, while the kinase/RNase domain alone remains strictly monomeric.

Why might the formation of oligomers rather than dimers be the key step in IRE1’s activation? The kinase/RNase domains can adopt two distinct dimeric conformations, known as the back-to-back and face-to-face orientations (*51*). The back-to-back orientation is thought to represent the RNase-active form of the protein, but is incapable of trans-autophosphorylation since both kinase active sites face outwards. Meanwhile, the face-to-face arrangement is perfectly suited for trans-autophosphorylation but is not expected to have RNase activity (*51*). It has been proposed that the dimerization of IRE1 allows the kinase/RNase domains to carry out trans-autophosphorylation in a face-to-face orientation, subsequently flipping around to form an active phosphorylated back-to-back dimer (*52*). However, this would require a massive rearrangement of cytosolic domains, and the feasibility of such a transition has not been demonstrated. Our data favor an alternative model (**Fig. 5**), wherein naïve IRE1 is either partially or fully pre-assembled into back-to-back dimers, which remain inactive due to their lack of phosphorylation. In response to ER stress, the lumenal domains drive the assembly of higher-order oligomers such as tetramers (that is, dimers of dimers). Since the back-to-back interfaces are already engaged, the formation of tetramers permits inter-dimer face-to-face interactions of kinase-RNase domains, thus enabling the phosphorylation reaction that in turn activates the RNase domains of the back-to-back dimers. Phosphorylation of the kinase domains of a dimer’s constituent protomers may occur either sequentially, via dissociation and reassociation of tetramers, or simultaneously, via the assembly of multiple dimers into a larger oligomer. The unimpeded ability of K599A to form dimers and oligomers further supports the notion that phosphorylation takes place at the level of higher-order oligomers rather than dimers.

Our results raise the question of whether the larger oligomers represent the maximally active form of IRE1 or are instead a phosphorylation-competent transient state *en route* to the generation of fully active phosphorylated dimers with and perhaps outside of oligomers. Our data showing the fast assembly and disassembly of high-order oligomers in response to saturating Tg, coupled with rapid hyper-phosphorylation, suggest that phosphorylated IRE1 may not need to remain oligomeric for its RNase to remain active. In fact, extensive phosphorylation either on the activation loop or elsewhere on IRE1’s kinase domain may serve as a negative feedback mechanism, as previously proposed for yeast IRE1 (*53*). A simple potential model for such negative feedback is that electrostatic repulsion across the face-to-face interface of two hyperphosphorylated dimers within an oligomer may break them apart into active back-to-back dimers and render the phosphate groups more accessible to the action of regulatory phosphatases (*46*).

Our data can be reconciled with seemingly contradictory earlier work to build a more comprehensive understanding of IRE1’s biology. First, the single-particle tracking approach does not rule out the existence of monomeric IRE1; it is entirely plausible that a small fraction of monomers remain in equilibrium with a largely dimeric resting population. Transient monomerization of the IF1^L^ interface, e.g. through the action of ER-lumenal chaperones (*25*), may play an important role in the regulated attenuation of IRE1 signaling in response to prolonged stress as previously suggested (*50*). Conversely, we do not claim that mammalian IRE1 lacks the capacity to assemble into clusters larger than tetramers; as with any oligomerization-prone protein, such clustering may largely be a function of protein expression level. IRE1 levels were found to be highly variable across a panel of cancer cell lines (*54*) and it is reasonable to suspect that large clusters of endogenous IRE1 do form in cell lines with elevated IRE1 expression. Rather than ruling out these possibilities, the present study demonstrates that IRE1 has the propensity to preassemble into inactive dimers in the absence of stress induction and that oligomerization past the tetrameric state is not strictly required for its RNase activation. In solution reactions, kinase/RNase dimers are capable of performing stem-loop endomotif-specific mRNA cleavage, while phosphorylated oligomers perform this function more efficiently than dimers and acquire a more promiscuous RNase activity termed RIDDLE (*16*). The present data suggest that in a cellular context, non-phosphorylated full-length IRE1 dimers are more restricted, perhaps by orientation, while even endomotif-directed RNase activity requires IRE1 oligomerization and phosphorylation. This restriction may be purely steric, due to the limited ability of the unphosphorylated membrane-tethered kinase/RNase dimers to adopt a conformation conducive to RNA cleavage, or it may arise from IRE1’s association with additional molecular players such as the ribosome (*55*) and/or the Sec61 translocon (*56*).

Protein oligomerization is a conceptually simple and common way in which information is communicated throughout the cell. Yet, experimental approaches for interrogating subtle oligomeric changes in the intracellular milieu remain scarce and fraught with caveats. We have developed an approach that is highly sensitive to the differences between monomers, dimers, and small oligomers in the plane of the ER membrane, while remaining easy to implement and, due to automated data analysis, resistant to researcher bias. Applying the approach to the key regulator of cellular proteostasis IRE1, we demonstrated that the dimer-to-oligomer transition serves as the primary regulatory step in IRE1 activation and reinforced the role of the lumenal domain as the master governor of IRE1’s oligomeric state. IRE1 has emerged as a highly promising molecular target in an ever-growing list of human diseases. Uncovering the basic principles behind its regulation promises to advance the design of future therapeutics, especially those intended to tune IRE1 activity through modulation of its oligomeric state.

## Materials and Methods

### Cell culture and experimental reagents

U-2 OS Flp-In T-REx cells were a kind gift of the Ivan Dikic lab and were independently authenticated through the human STR profiling service offered by the American Type Culture Collection (ATCC). Cells were cultured in high-glucose DMEM (Thermo Fisher) supplemented with 10% tetracycline-free fetal bovine serum (FBS; Takara Bio), 6 mM L-glutamine, and 100 U/ml penicillin/streptomycin. All cell lines used in the study tested negative for mycoplasma contamination when assayed with either the Universal Mycoplasma Detection Kit (ATCC 30-1012K) or the MycoAlert Detection Kit (Lonza LT07-418). Tunicamycin and thapsigargin were purchased from Sigma-Aldrich or from Tocris. JF549 and JF646 dyes conjugated with the HaloTag ligand were a kind gift of Luke Lavis (Janelia Farms). The antibodies used for immunoblotting are listed in the *Immunoblotting* section.

### Endogenous tagging of IRE1 in U-2 OS cells

To achieve full editing despite the hyperploid nature of the U-2 OS cells, we first generated a partial IRE1α knockout cell line harboring a single intact allele of *ERN1*, the gene encoding IRE1α (cell line ID: PWM359). This was done using the same CRISRP/Cas9-based approach that we used to generate a complete IRE1α knockout in our previous paper (*21*), except that rather than looking for a clone that contained no copies of WT *ERN1*, we identified clones that contained a single unedited allele. The presence of a single intact *ERN1* allele was confirmed by TOPO cloning and immunoblotting. These partial knock-out cells were then co-transfected with a plasmid encoding Cas9 with the guide RNA and a homology-directed repair (HDR) template plasmid targeted at C-terminus of *ERN1*. Design of both plasmids followed the protocol published elsewhere (*57*). Edited cells were selected by fluorescence-activated cell sorting (FACS), separated into clonal populations by limiting dilution, and assayed for IRE1α expression and UPR activation by immunoblotting and RT-PCR. Two clones were selected for further study: a somewhat higher expressing clone (cell line ID: PWM360) and a somewhat lower expressing clone (cell line ID: PWM361). When immunoblotted against IRE1α, both clones produced a clear band that ran slower than WT IRE1α, indicating a successful integration of the full-length HaloTag.

### Sample preparation for microscopy

Cells were seeded at a density of 1.6 x 10^4^ cells/cm^2^ into glass-bottom 8 well chamber slides (ibidi 80827), which were pre-coated with rat tail collagen type I (Corning 354236) at 10 μg/cm^2^ in accordance with the manufacturer’s instructions (briefly, a 2-hour incubation at room temperature). Twenty-four hours prior to imaging, the growth medium was replaced with “Imaging medium”: FluoroBrite DMEM (ThermoFisher) supplemented with 10% tetracycline-free fetal bovine serum (FBS; Takara Bio) and 6 mM L-glutamine, without antibiotics. For experiments requiring transfection, cells were transfected with a mixture of 50 ng of plasmid DNA and 50 ng of carrier salmon sperm DNA per well immediately following medium change (i.e. 24 hours prior to the start of imaging). Transfections were carried out in “Imaging medium” using the Fugene HD transfection reagent (Promega).

On the day of imaging, cells were treated with ER stressors at the indicated time points. Labeling with JF549 and JF646 dyes conjugated with the HaloTag ligand was initiated 1.5 hours prior to the start of imaging. First, the dyes were added to pre-warmed “Imaging medium”, and this medium was used to replace the cells’ growth medium. We experimentally found the optimal molar dye ratio to achieve ~50% labeling with each ligand to be 1:20 (5 nM JF549-HaloTag and 100 nM JF646-HaloTag). The large difference in required concentrations is likely due to the difference in membrane permeability between the two dyes. We experimentally found the 5 nM JF549 / 100 nM JF646 concentrations to be saturating under our labeling conditions since further increases in dye amounts did not lead to a further increase in the density of diffusing spots in IRE1-HaloTag cells. Following addition of the medium containing the two dyes (and any required ER stressors), cells were returned to the incubator for 1 hour. Then, cells were washed twice with warm PBS, washed once with pre-warmed “Imaging medium”, and returned to the incubator for an additional 5 minutes to give any unbound dye time to diffuse out of the cells. The medium was replaced one more time with pre-warmed “Imaging medium” containing any required ER stressors to finish sample preparation.

### Microscopy

All imaging was carried out on one of two Nikon Ti-E inverted microscopes (#1 and #2 hereafter), each equipped with a Nikon motorized TIRF module, an Agilent/Keysight MLC400 fiber-coupled laser light source, a Perfect Focus System (PFS, Nikon), a 100x 1.49 NA oil immersion objective (Apo TIRF, Nikon), and a Hamamatsu Flash 4.0 CMOS camera. Microscope #1 held a ZET405/488/561/640m-TRFv2 quadruple bandpass filter cube (Chroma), while microscope #2 held a ZET488/561/640m triple bandpass filter cube (Chroma). Additionally, microscope #1 included a Yokogawa CSU-X high-speed confocal scanner unit and an Andor iXon 512 × 512 EMCCD camera, which were used for spinning-disk confocal microscopy experiments. Both microscopes featured full temperature and CO_2_ control to maintain the samples at 37°C and 5% CO_2_, one using a custom-built enclosure (#1) and the other using an OkoLab Live stage insert (#2). All components of microscope #1 were controlled by the μManager open-source platform (*58*), while microscope #2 was controlled with NIS-Elements software (Nikon).

Oblique-angle illumination conditions were achieved by focusing on a cell, engaging the PFS, and gradually increasing illumination angle with the motorized TIRF lens until single-molecule spots along the bottom surface of the cell became clearly visible. Videos were acquired with a 60 ms combined frame time, split into a 25 ms exposure in the JF549 channel (561 nm laser, operated at 25 mW) and a 25 ms exposure in the JF646 channel (640 nm laser, operated at 40 mW), with the remaining 10 ms accounting for channel switching times. The two channels were imaged sequentially by the same camera using camera-triggered switching of the acousto-optic tunable filter (AOTF) built into the light source. Frames were cropped to approximately 500×500 pixels prior to acquisition since the full camera sensor could not be read out fast enough to support the required frame rate. Typically, 100 combined frames were acquired per cell (6 second total movie duration), which in our hands provided a good number of trajectories per cell while avoiding extensive photobleaching of the dyes. To locate and choose cells for imaging, we used the full size of the camera sensor and acquired a series of tiled snapshots of an area containing ~100 cells. We then selected cells that were morphologically normal, well-adhered, and spread out. When imaging transiently transfected cells, we chose cells in which the HaloTag-labeled proteins were expressed at sufficiently low levels to allow us to clearly see individual spots corresponding to single molecules.

### Data analysis

Single-molecule data were analyzed to identify co-localizing two-color trajectories using a pipeline developed in house, described in detail below. First, each movie was split into the two individual channels, JF549 and JF646. Next, spots were located using the Laplacian of Gaussian (LoG) detector implemented in the TrackMate plugin (*59*) for ImageJ. Then, identified spots were tracked using the Linear Assignment Problem (LAP) algorithm (*60*), also implemented within the TrackMate plugin. All input parameters for both the LoG detector and the LAP tracker were chosen empirically to match the expected output in a subset of randomly selected single-molecule movies; afterwards, they were kept constant for the analysis of all data used to construct the plots presented in this paper. To speed up analysis and ensure that the exact same settings are used to process every movie, we scripted TrackMate to read all settings from a standardized JSON configuration file and perform both spot detection and tracking on all movie files contained within a given folder. The TrackMate output files containing spot and track data for each channel were then saved to disk in the XML format for further analysis.

All subsequent analysis was performed in Python. The broad goal of this analysis was to identify tracks that correlated well in space and time between the JF549 and JF646 channels. To avoid problems imposed by uneven track durations and trajectories crossing each other, we decided to perform the analysis using a short sliding window. In other words, instead of considering the entire movie at once, we binned each movie into overlapping shorter movies containing a fixed number of frames each, and looked for co-localizing trajectories in each of the shorter movies. To achieve this, the TrackMate output files were parsed and filtered to only include tracks that span at least as many frames as the length of the sliding window. The sliding window was then moved across the duration of the movie in 1 frame increments. In each of the resulting windows, only tracks that were fully defined within that window (i.e., had position information for each frame) were selected for correlation analysis.

Pearson’s correlation coefficients (PCC) were then individually calculated for the X- and Y-coordinates of every pair of spatially adjacent tracks (adjacent meaning that at least a subset of data points of track B are contained within the rectangle that bounds track A). The requirement for tracks being spatially adjacent both increased the computational efficiency of the algorithm and helped eliminate false positives from short tracks of similar shapes that occurred by chance in different parts of the cell. A pair of tracks was determined to be correlated if all of the following conditions were met: 1) the two tracks share at least one window *N* frames long in which the position of each spot is well-defined in every frame, 2) At least one such window yields a PCC value greater than *T* for both the X- and Y- coordinates of the tracked spot, and 3) The two tracks are at least partially overlapping in space. The value plotted in the figures, “% correlated trajectories”, is defined as follows:

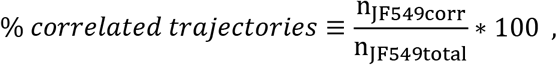

where n_JF549total_ is the total number of trajectories in the JF549 channel that are at least as long as the length of the sliding window (*N*), while n_JF549corr_ is the number of trajectories in the JF549 channel that are found to have correlated trajectories in the JF646 channel as described above.

In our analysis, the only user-selected parameters that tune the sensitivity of the approach are *N* (the length of the window, in frames) and *T* (the threshold value for the PCC). Just as with the tracking algorithm, we first empirically found values of *N* and *T* that yielded robust identification of visually correlated tracks without giving too many false positives, and then used these values in all subsequent data analysis. We did find that due to differences in filters, laser intensity, and alignment between the two microscopes, a different combination of *N* and *T* yielded the highest dynamic range in our assay. Data from each microscope were fully internally consistent but we avoided showing data collected on two different microscopes on the same plot. Thus, each panel in the paper contains either data collected exclusively on microscope #1 or on microscope #2.

To speed up data processing and enhance reproducibility, we again scripted the analysis to read a single JSON configuration file that specifies the *N* and *T* parameters, along with a full list of folders containing the TrackMate XML files for each condition. The code reports the fraction of correlated tracks for each condition with 95% confidence intervals determined by bootstrapping. The README.md file included with the source code (*61*) contains detailed instructions for running this analysis and replicating all plots in the paper from source data (*62*, *63*). In organizing the analysis software, we sought to make reproducing our data and adapting the code to different single-molecule co-localization studies as straightforward as possible.

### Estimation of IRE1 cluster stoichiometry

A simple yet useful model for estimating cluster stoichiometry based on the fraction of correlated tracks works as follows. Assume that each HaloTag-conjugated protein can occupy one of three states: bound to an unbleached JF549 dye molecule, bound to an unbleached JF646 dye molecule, or undetectable. The latter category is a catch-all for every possible reason a protein may escape detection such as dye bleaching, incomplete labeling, new protein synthesis after labeling reaction, and false negatives in the spot detection algorithm. Let the probabilities of these three states be denoted as *P*_*1*_ (JF549-bound), *P*_*2*_ (JF646-bound) and *P*_*u*_ (undetectable). Because the combined probabilities must add up to unity, *P*_*u*_ = *1* – *P*_*1*_ – *P*_*2*_. Then, for a cluster comprised of *n* individual molecules, we can express the total probability that the cluster contains at least one dye of each color as follows:

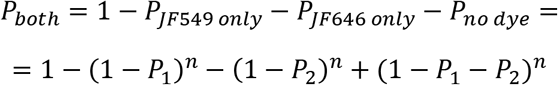

However, in our experiment, the clusters containing no detectable dyes are invisible, and what we measure experimentally is instead the observed fraction of all visible clusters that contain at least one dye of each color. Let’s call this quantity the fraction of observed co-localizers, *F*_*obs*_:

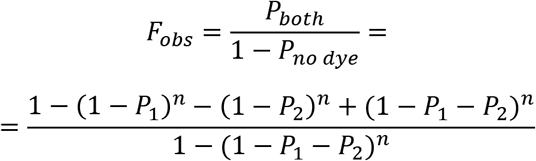

To estimate cluster stoichiometry based on the experimentally measurable F_obs_, we first need a measurement of P_1_ and P_2_, which can be done using data from the constitutive 2x HaloTag homodimer construct, where we know that *n =* 2. Let’s assume that *P*_*1*_ = *P*_*2*_ = *P*_*L*_ (labeling probability), since all our experiments are done in a regime where the labeling densities with the two different dyes are nearly identical. Plugging these assumptions into the expression for *F*_*obs*_, we obtain:

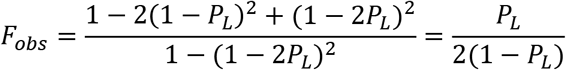

Rearranging this expression, we find that P_L_ can be expressed in terms of *F*_*obs*_:

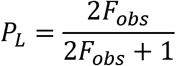

Once we have an estimate of *P*_*L*_ from the 2x homodimer control (in our experiment, this value is typically around 0.14), it can be simply plugged into the earlier expression for *F*_*obs*_ and plotted as a function of *n* to yield an estimate of average cluster stoichiometry for any value of *F*_*obs*_. Of course, this model is a significant oversimplification of the true underlying processes (mainly due to lumping all possible sources of error into the single term *P*_*u*_), but it does provide a useful ballpark estimate.

### XBP1 mRNA splicing assays

Cells were grown in wells of a 12-well plate, treated with ER stressors as indicated in the figure, and harvested at ~70% confluency with TRIzol (Thermo Fisher) in accordance with the manufacturer’s instructions. RNA was then extracted from the aqueous phase using a spin column-based purification kit (RNA Clean & Concentrator-5, Zymo Research # R1015) and reverse-transcribed into cDNA using SuperScript VILO Master Mix (Thermo Fisher # 11755050). The cDNA was diluted 1:10 and used as a template for PCR with the following primer pair: VB_pr259 (CGGAAGCCAAGGGGAATGAA) and VB_pr167 (ACTGGGTCCAAGTTGTCCAG). PCR was carried out with Taq polymerase (Thermo Fisher # 10342020) in the manufacturer-supplied Taq buffer supplemented with 1.5 μM Mg^2+^. The following PCR program was used: (1) Initial denaturation: 95°C for 2 minutes, (2) 95°C for 30 seconds, (3) 60°C for 30 seconds, (4) 72°C at 30 seconds, (5) Repeat steps 2-4 27 more times, for 28 total PCR cycles. PCR products were visualized on a 3% agarose gel stained with SYBR Safe (Thermo Fisher S33102) and imaged on a ChemiDoc gel imaging system (BioRad).

### Immunoblotting

Cells were grown in 6-well plates in RPMI1640 or DMEM media supplemented with 10% (v/v) FBS (Sigma), 2 mM glutamine (Gibco), and 100 U/mL penicillin plus 100 μg/mL streptomycin (Gibco), and treated as indicated. Thapsigargin (Tocris) was used at a concentration of 100 nM and tunicamycin (Tocris) at 5 μg/mL, dissolved in DMSO. DMSO was used as the untreated control.

Cells were trypsinized using Trypsin-EDTA 0.05% and protein lysates were extracted in RIPA buffer (EMD Millipore) with Halt protease and phosphatase inhibitor cocktail (Thermo Scientific). The crude lysates were cleared by centrifugation at 13,000 rpm for 15 min and protein content was analyzed by Pierce BCA protein assay (Thermo Scientific).

Equal amounts of protein (40 μg/condition) were run with SDS-PAGE and electrotransferred onto membranes that were blocked with 5% dried nonfat milk powder in TBST (blocking solution). Blots were incubated with 1/1000 dilution in 5% blocking solution of primary antibodies overnight at 4C. Antibodies (Abs) for IRE1α (3294), PERK (3192), ATF4 (11815), CHOP (2895) were from Cell Signaling Technology (CST). ATF6 antibody (66563-1) was from Proteintech. β-actin (5125) from CST was used as a housekeeping control. Abs for XBP1s and pIRE1 were generated at Genentech and have been described elsewhere(*46*). Blots were washed in TBST, then incubated during 1h at room temperature with 1/10,000 dilution of the corresponding peroxidase-conjugated secondary antibodies in blocking solution: donkey anti-rabbit and anti-mouse from Jackson Immunoresearch. Blots were finally washed in TBST and analyzed using Super Signal West Dura or Femto (Thermo Scientific).

## Availability of materials, data, and software

The code used to analyze raw data and generate all figures in this paper is freely available from Zenodo (*61*). All raw single-molecule microscopy data are available from Dryad (*62*). All other raw data, including full gel images, together with processed single-molecule microscopy data, are available from Zenodo (*63*). Additionally, raw data and code are backed up on the Walter Lab server and are available upon request in case there is an issue with one of the databases listed above. All cell lines and constructs used in this paper are available upon request.

## Acknowledgements

We thank members of the P.W. and A. Ashkenazi laboratories for helpful discussions, especially Smriti Sangwan, Tsan-wen Lu, Adrien Le Thomas, David A. Lawrence, Silvia Ramundo, and Morgane Boone. We thank Nico Stuurman for advice and help with single-molecule microscopy and Elif Karagöz (Max Perutz Labs, Universität Wien, Vienna, Austria) for comments on the manuscript. We thank David Ron (Cambridge Institute for Medical Research, University of Cambridge, Cambridge, United Kingdom) for proposing the IRE1-GST dimerization control in Supplementary Figure 1. We thank Ivan Dikic (Institute of Biochemistry II, School of Medicine, Goethe University, Frankfurt am Main, Germany) and Luke Lavis (Janelia Research Campus, Howard Hughes Medical Institute, Ashburn, VA, USA), for reagents and advice.

## Author contributions

V.B., I.Z.-G., A. Alamban, A. Ashkenazi, and P.W. designed research; V.B., I.Z.-G., and A. Alamban performed research; V.B. contributed new reagents and analytic tools; V.B., I.Z.-G., A. Ashkenazi and P.W. wrote the paper.

