## Supplemental Information for "Endoplasmic reticulum stress activates human IRE1α through reversible assembly of inactive dimers into small oligomers"

### Supplementary information

**Supplementary Table 1: All plasmids used in this study.**

| Plasmid ID | Plasmid name | Resistance | Description |
|---|---|---|---|
| pPW3754 | SpCas9 and gRNA targeting the C-terminus of HsIRE11 | Ampicillin | Expression of human codon-optimized SpCas9 and gRNA targeting the C-terminus of human IRE1. |
| pPW3755 | HDR-HsIRE1a-10xGS-HaloTag | Ampicillin + G418 (mammalian ) | Complete HDR template for cloning a C-terminal 10xGS-HaloTag into human IRE1alpha. Should be co-transfected with a plasmid encoding the corresponding gRNA and Cas9. Contains a mutated PAM site to ensure that the genome is no longer cut after a successful HDR event. |
| pPW3756 | CMVd3-ERmembrane-HaloTag-KKMP | Kanamycin | Construct for low-level transient expression of a single HaloTag protein targeted to the ER membrane. ER targeting is achieved by an IRE1-derived signal peptide and TM helix, with a C-terminal KKMP ER retention signal. Expression is driven by the heavily truncated CMVd3 promoter. |
| pPW3757 | CMVd3-ERmembrane-2xHaloTag-KKMP | Kanamycin | Construct for low-level transient expression of two tandem HaloTag proteins targeted to the ER membrane. |
| pPW3758 | CMVd3-HsIRE1-HaloTag | Kanamycin | Construct for low-level transient expression of full-length HsIRE1 with a C-terminal HaloTag, with the exact same 10x GS linker sequence as that in pPW3755. Expression is driven by the heavily truncated CMVd3 promoter. |
| pPW3759 | CMVd3-HsIRE1deltaLD-HaloTag | Kanamycin | Construct for low-level transient expression of delta-lumenal domain HsIRE1 with a C-terminal HaloTag. |
| pPW3760 | CMVd3-HsIRE1-K599A_KinaseDead-HaloTag | Kanamycin | Construct for low-level transient expression of kinase-dead (K599A) HsIRE1 with a C-terminal HaloTag. |
| pPW3761 | CMVd3-HsIRE1(WLLI-GSGS)[359-362]-HaloTag | Kanamycin | Construct for low-level transient expression of HsIRE1 with a lumenal domain IF2 mutation (WLLI-GSGS) [359-362] with a C-terminal HaloTag. |
| pPW3762 | CMVd3-HsIRE1(K121Y)-HaloTag | Kanamycin | Construct for low-level transient expression of HsIRE1 with a lumenal domain IF1 mutation (K121Y) with a C-terminal HaloTag. |
| pPW3763 | CMVd3-HsIRE1dLKR-HaloTag | Kanamycin | Construct for low-level transient expression of delta-LKR HsIRE1 with a C-terminal HaloTag. |
| pPW3781 | CMVd3-ERmembrane-GST-HaloTag-KKMP | Kanamycin | Construct for low-level transient expression of a single GST-fused HaloTag protein targeted to the ER membrane. The GST fusion causes this to be a constitutive dimer. |

**Supplementary Table 2: All cell lines used in this study.**

| Cell line ID | Cell line name | Origin | Description |
|---|---|---|---|
| PWM253 | U-2 OS WT T-REx Flp-In | DOI: 10.1073/pnas.1915311117 | Parental cell line for all cells in this study. Kind gift of Ivan Dikic. |
| PWM254 | U-2 OS IRE1α KO | DOI: 10.1073/pnas.1915311117 | CRISPR knock-out of IRE1α in PWM253 (see reference for details) |
| PWM359 | U-2 OS IRE1α partial KO | Generated for this study | Partial CRISPR knock-out of IRE1α in PWM253, containing one intact ERN1 allele |
| PWM360 | U-2 OS IRE1a-HaloTag endogenously tagged | Generated for this study | Introduction of a C-terminal HaloTag into the endogenous ERN1 locus of PWM359 cells; clonal population. |
| PWM361 | U-2 OS IRE1a-HaloTag endogenously tagged, low expression clone | Generated for this study | Introduction of a C-terminal HaloTag into the endogenous ERN1 locus of PWM359 cells; clonal population with lower IRE1 expression level than PWM360. |

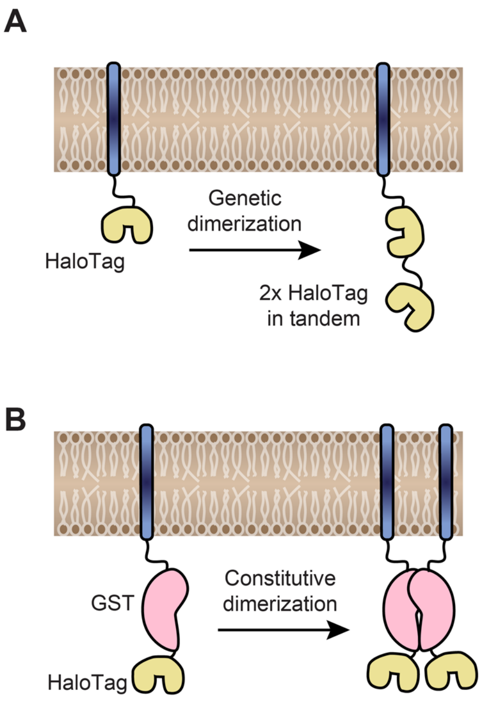


**Supplementary Figure 1: Orthogonal dimerization controls**. (A) Schematic representation of the genetic dimerization strategy used as a primary control throughout the manuscript. A dual-HaloTag construct is engineered by fusing two HaloTag proteins in tandem, separated by a flexible GS linker, to the C-terminus of a single transmembrane helix targeted to the ER membrane. (B) Orthogonal dimerization control used to rule out the possibility that the internal and C-terminal HaloTags of the construct shown in panel A may have different labeling efficiencies. In this control, ER membrane-tethered HaloTag proteins assemble into constitutive dimers via an internal GST tag. (C) Single-particle tracking results comparing the fraction of correlated trajectories for the constructs shown in panels A and B. Each data point represents a single cell. Error bars represent 95% confidence intervals.

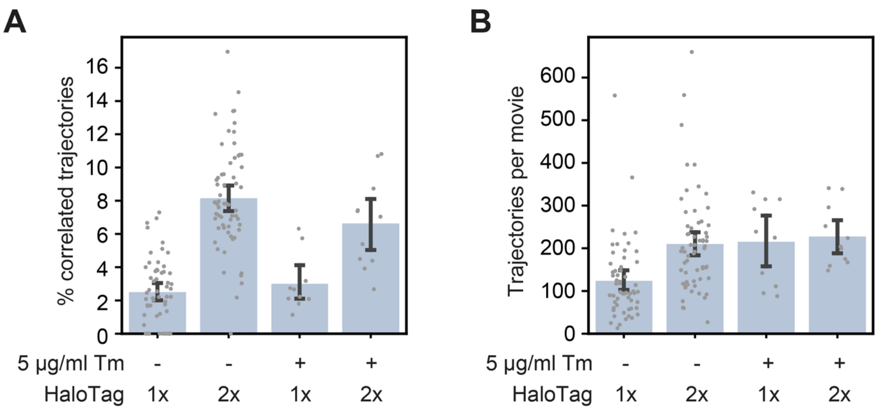


**Supplementary Figure 2: Effect of ER stress on HaloTag controls**. (A) Single-particle tracking data showing the fraction of correlated trajectories for the 1x and 2x HaloTag controls, with and without a 4-hour treatment with tunicamycin. (B) Number of trajectories per movie for the four conditions shown in panel A, demonstrating that the changes in % correlated trajectories are independent of construct expression levels. Each data point represents a single cell. Error bars represent 95% confidence intervals.

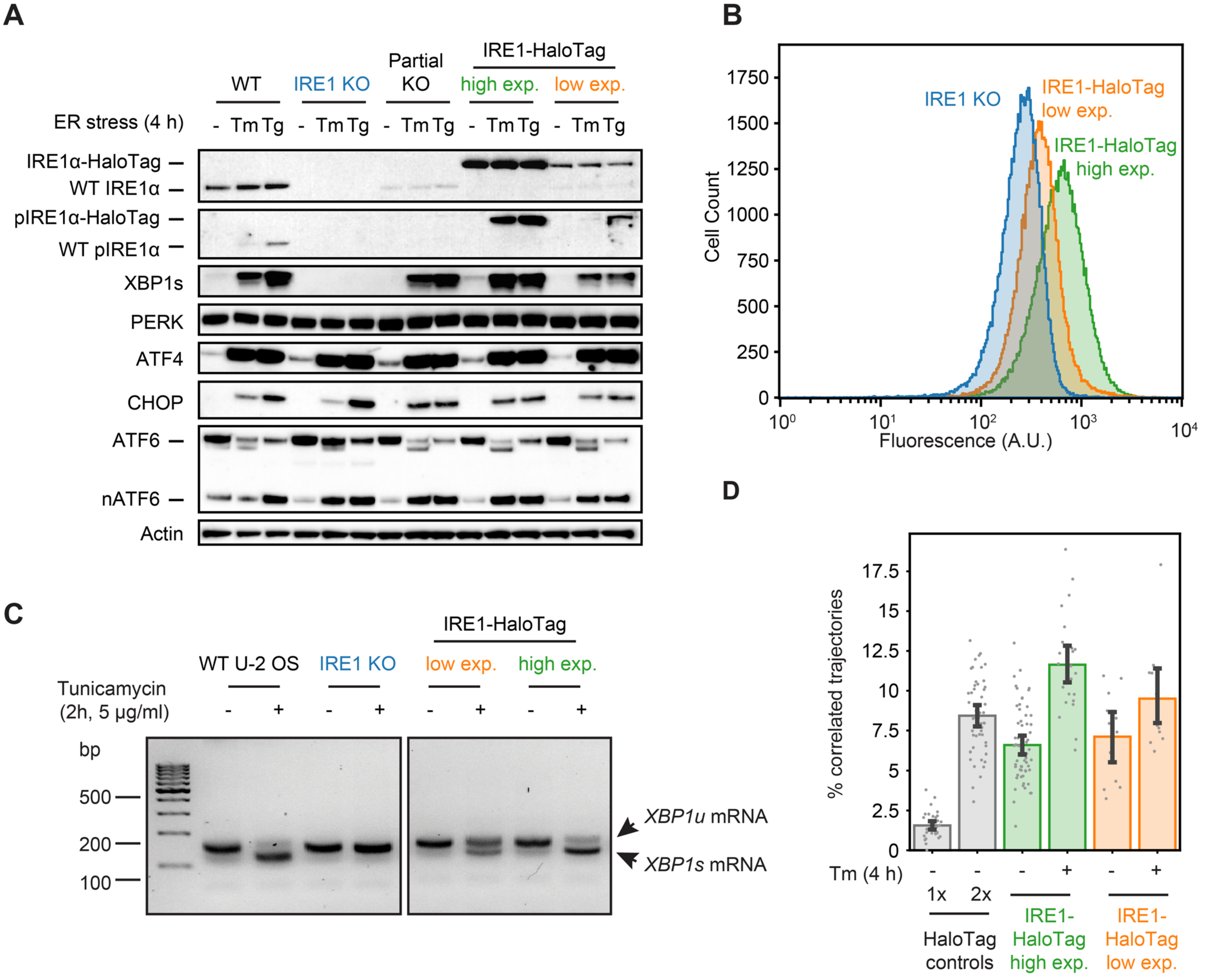


**Supplementary Figure 3: Comparison of high- and low-expression clones**. (A) Immunoblot showing IRE1 expression levels and UPR activation in WT U-2 OS cells, IRE1 KO U-2 OS cells, partial KO cells used as the parental cell line for generating HaloTag knock-ins, and two clones of endogenously labeled HaloTag (with high and low IRE1 expression levels). Note the large shift in protein size due to the addition of the HaloTag and the absence of a WT IRE1 band in the two clones on the right. (B) Flow cytometry analysis of the low-and high-expressing clones shown in panel A. Cells were labeled with 5 nM JF549-HaloTag dye for 1 hour prior to the start of the flow cytometry experiment. Note the unimodal intensity distributions of both clones, ruling out the possibility that clone PWM361 simply contains a bimodal mixture of low- and high-expressing cells. (C) RT-PCR analysis of *XBP1* mRNA splicing by the clones shown in panels A and B. (D) Single-particle tracking data showing stress-dependent oligomerization of the clones shown in the previous panels. IRE1 in the lower-expressing clone remains dimeric in unstressed cells, while the shift to higher-order oligomers upon stress is less prominent than in the higher-expressing clone. Each data point represents a single cell. Error bars represent 95% confidence intervals.

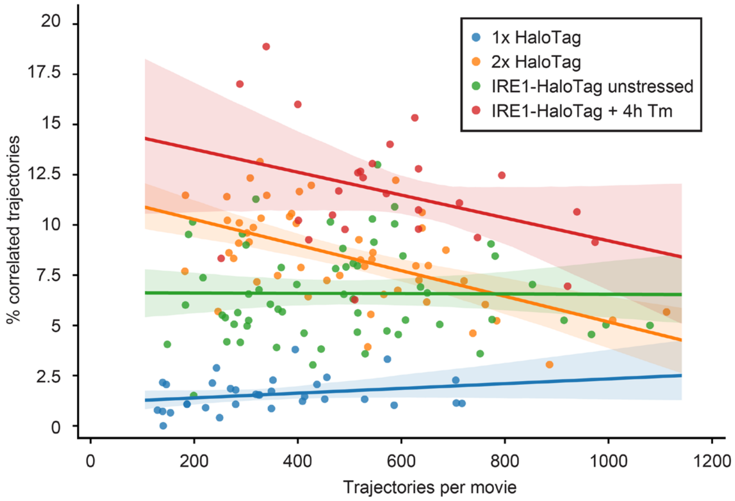


**Supplementary Figure 4: Trajectory density vs. percent correlation**. A plot showing the relationship between the percentage of a calculated trajectories in a given cell against the number of trajectories in the movie collected from that cell, for the four conditions indicated in the box. Solid lines represent linear fits, with the shaded regions around them showing 95% confidence intervals. Each data point represents a single cell.

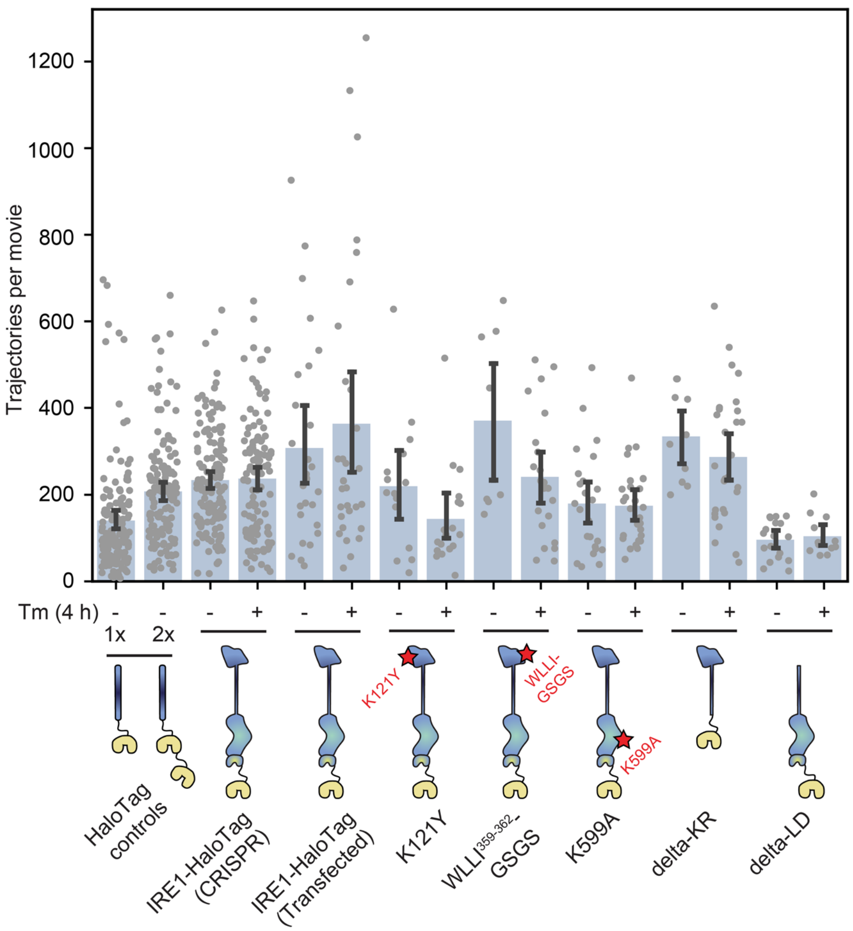


**Supplementary Figure 5: Trajectory counts of all mutants**. Number of trajectories per movie for all mutants shown in Figure 5, demonstrating that all constructs are expressed at comparable levels and that the changes in % correlated trajectories are independent of construct expression levels. Each data point represents a single cell. Error bars represent 95% confidence intervals.

**Supplementary Movie 1: Co-localizing spots in cells expressing 2x tandem HaloTag.** A cropped and annotated movie recorded from an IRE1 KO cells transiently transfected with the 2x tandem HaloTag construct and labeled with a mixture of JF549 and JF646 dyes. Two separate co-localizing spots (as determined by the automated analysis pipeline) are annotated.
